# Can static optimization detect changes in peak medial knee contact forces induced by gait modifications?

**DOI:** 10.1101/2022.07.18.500342

**Authors:** Janelle M. Kaneda, Kirsten A. Seagers, Scott D. Uhlrich, Julie A. Kolesar, Kevin A. Thomas, Scott L. Delp

**Author notes:** Please direct correspondence to: Janelle M. Kaneda.

## Abstract

Medial knee contact force (MCF) is related to the pathomechanics of medial knee osteoarthritis. However, MCF cannot be directly measured in the native knee, making it difficult for therapeutic gait modifications to target this metric. Static optimization, a musculoskeletal simulation technique, can estimate MCF, but there has been little work validating its ability to detect changes in MCF induced by gait modifications. In this study, we quantified the error in MCF estimates from static optimization compared to measurements from instrumented knee replacements during normal walking and seven different gait modifications. We then identified minimum magnitudes of simulated MCF changes for which static optimization correctly identified the direction of change at least 70% of the time. A full-body musculoskeletal model with a multi-compartment knee and a custom static optimization implementation was used to estimate MCF. Simulations were evaluated using experimental data from three subjects for a total of 115 steps. Static optimization underpredicted the first peak (mean absolute error = 0.16 bodyweights) and overpredicted the second peak (mean absolute error = 0.31 bodyweights) of MCF. Average root mean square error in MCF over stance phase was 0.32 bodyweights. Static optimization detected the direction of change with at least 70% accuracy for early-stance reductions, late-stance reductions, and early-stance increases in peak MCF of 0.10 bodyweights and greater. These results suggest that a simple static optimization approach accurately detects directional effects on early-stance medial knee loading, potentially making it a valuable tool for evaluating the biomechanical efficacy of gait modifications for knee osteoarthritis.

## Introduction

Knee osteoarthritis is a leading cause of disability (Vos et al., 2012) and affects over 654 million individuals worldwide (Cui et al., 2020). The disease is often isolated to the medial compartment, which has 5-10 times higher prevalence compared to lateral knee osteoarthritis (Ahlbäck, 1968; Felson et al., 2002; Jones et al., 2013). Andriacchi and colleagues (2004) have suggested that medial knee osteoarthritis is associated with compressive medial loading. The knee adduction moment is an easier-to-estimate surrogate for compressive medial knee contact force (MCF), and relates to medial knee osteoarthritis progression (Miyazaki et al., 2002) and severity (Sharma et al., 1998). Therefore, gait modifications have aimed to reduce the peak knee adduction moment (Fregly et al., 2007; Shull et al., 2013; Simic et al., 2011). Recently, MCF has been related to cartilage loss, and thus medial knee osteoarthritis progression (Brisson et al., 2021). Since changes in the knee adduction moment do not fully describe changes in MCF (Meyer et al., 2013b; Richards et al., 2018; Walter et al., 2010), load-reducing interventions may be more effective if they aim to reduce MCF (Kinney et al., 2013b).

Musculoskeletal simulation is often used to estimate MCF. Although there are various methods to estimate joint loading (Lloyd and Besier, 2003; Pizzolato et al., 2015), static optimization is a simple and commonly used technique. This method solves for muscle forces by optimizing muscle activations based on ground reaction force and motion data. It is important to test the accuracy of simulation estimates of MCF against measurements from instrumented knee replacements (D’Lima et al., 2012; Hicks et al., 2015). In the context of gait modifications, it is critical to evaluate directional accuracy, which we define as the ability to predict an increase or decrease in MCF induced by a gait modification compared to natural walking (Kinney et al., 2013b). Previous studies have reported MCF and total knee contact force errors during stance phase using static optimization (Brandon et al., 2014; DeMers et al., 2014; Lerner et al., 2015; Lundberg et al., 2013), but there has been less work testing the ability of static optimization to detect changes in peak MCF across multiple individuals and gait modifications.

To address this gap in knowledge, we sought to: (1) quantify the errors in MCF estimated from static optimization and (2) identify when static optimization can accurately determine whether a gait modification increases or decreases MCF. All simulations used in this study are freely available at https://simtk.org/projects/statop_val.

## Methods

### Overview

We simulated knee contact force using a musculoskeletal model and a custom static optimization implementation for three individuals and seven gait modifications. To compare the simulated and experimental contact forces, we evaluated mean absolute error at peaks of contact force and root mean square error over stance phase. We also determined the accuracy of correctly predicting the direction of change in peak contact force induced by gait modifications (i.e., directional accuracy).

### Experimental data

We analyzed data from three unique subjects in the 3^rd^, 4^th^, and 6^th^ Grand Challenge competitions for predicting knee loads (Fregly et al., 2012; Kinney et al., 2013a), including instrumented knee forces, motion capture, and force plate data from overground gait trials for natural walking (baseline) and seven different gait modifications (Table 1). We added virtual hip joint centers estimated from thigh and pelvis kinematics during flexion-extension and abduction-adduction trials (Fregly et al., 2012; Piazza et al., 2004).

**Table 1.** Demographics and number of steps per walking pattern evaluated for three Grand Challenge subjects.

|  | 3<sup>rd</sup> Grand Challenge | 4<sup>th</sup> Grand Challenge | 6<sup>th</sup> Grand Challenge |
| --- | --- | --- | --- |
| **Demographics (Fregly et al., 2012)** |  |  |  |
| **Leg alignment** | Neutral | Neutral | Valgus |
| **Body mass index (kg/m<sup>2</sup>)** | 28.1 | 23.6 | 23.7 |
| **Gender** | Female | Male | Male |
| **Number of steps per walking pattern** |  |  |  |
| **Baseline** | 5 | 5 | 4 |
| **Bouncy** | 4 | 6 | 6 |
| **Crouch** | 4 | Mild (6) | 5 |
| **Forefoot strike** | 5 | 6 | 4 |
| **Medial thrust** | 5 | 8 | N.A.* |
| **Smooth** | 5 | N.A.* | 2 |
| **Trunk sway** | 4 | N.A.* | N.A.* |
| **Walking poles** | Long (2)<br>Short (3) | Long narrow (6)<br>Long wide (6)<br>Short narrow (7)<br>Short wide (7) | N.A.* |

\* Not Available. The given walking pattern was not included in the dataset for the given subject.

Instrumented knee replacements measured total contact force, and we calculated MCF and lateral contact force using regression equations (3^rd^ Grand Challenge: Fregly et al., 2012; 4^th^ Grand Challenge: Zhao et al., 2007; 6^th^ Grand Challenge: Meyer et al., 2013a). Simulated and measured knee contact forces were low-pass filtered at 15 Hz (zero-lag, 4^th^ order Butterworth). We removed trials with erroneous experimental marker, ground reaction force, and instrumented knee data.

We defined the first and second contact force peaks as the maximum value between 15-35% and 65-85% of stance, respectively. We selected these ranges as ±10% around 25% and 75% of stance, which are commonly used for defining early-stance and late-stance peaks (Fregly et al., 2009).

### Musculoskeletal simulation

We integrated a multi-compartment knee (Lerner et al., 2015) into a full-body model (Rajagopal et al., 2016) to compute medial and lateral compartment contact forces. Using the Scaling Tool in OpenSim 4.0 (Seth et al., 2018), we used standing static calibration poses to scale the model. We used the OpenSim Inverse Kinematics Tool to solve for the kinematics that minimized the sum-squared error between experimental and virtual markers.

We used a custom static optimization implementation in MATLAB R2017b (Mathworks Inc., Natick, MA, USA) utilizing OpenSim, which includes passive muscle forces and tendon compliance (Uhlrich et al., 2022). The objective function minimized the sum of squared muscle activations (Anderson and Pandy, 2001). We used the OpenSim Joint Reaction Analysis Tool to estimate MCF and lateral contact force from static optimization muscle force estimates; total contact force was the sum of these compartmental loads. Further simulation details are described in Supplementary Note 1.

### Error evaluation

We compared simulated and measured knee contact forces to evaluate the simulation results (Fig. 1, Supplementary Fig. 1, Supplementary Table 1, Supplementary Note 2). We calculated mean absolute error at the two peaks of MCF during the stance phase for all trials of baseline and gait modifications (*N*=115 trials), and also for changes from baseline (*N*=101 gait modification trials; Supplementary Table 2). We calculated root mean square error over stance phase and averaged over all trials.

**Figure 1.**
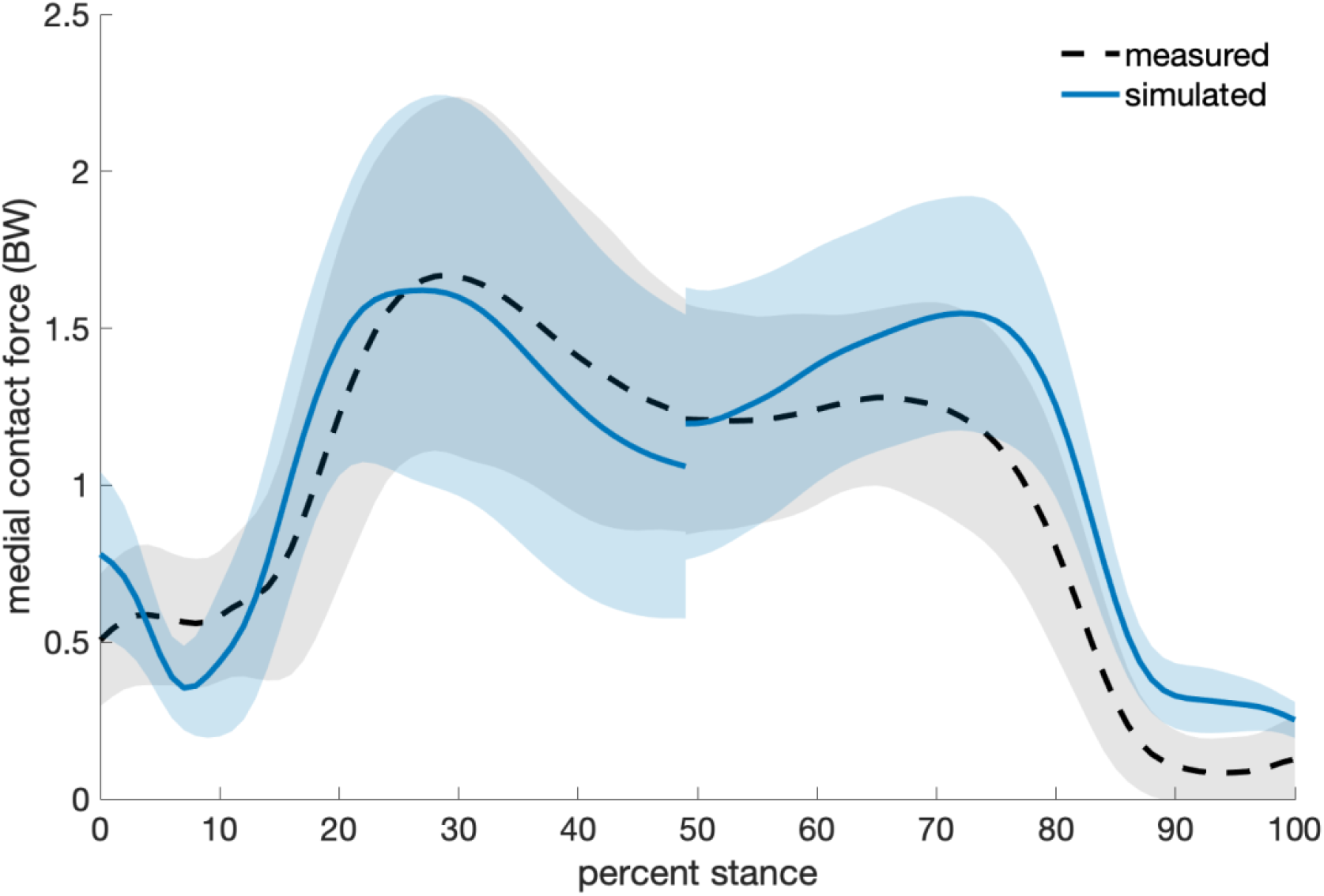
Mean ± standard deviation (shaded) of measured and simulated medial knee contact force in bodyweights (BW) for all evaluated trials, including natural walking (#=115). The discontinuity at 50% of stance results from computing early-stance and late-stance contact forces from different gait cycles for some steps.

It is important to identify the magnitudes of change in contact force that static optimization can accurately detect. Thus, we calculated directional accuracy for various thresholds of simulated knee contact force changes to identify the minimum magnitudes of change that represented true increases or reductions (Fig. 2, Supplementary Fig. 2, Supplementary Table 3, Supplementary Note 2). We calculated directional accuracy by comparing the directional changes from simulations and experiments. We calculated directional change by subtracting the subject’s average baseline peak from each gait modification trial peak, in both early and late stance. Thus, positive changes represent increases from baseline, and negative changes represent decreases from baseline.

**Figure 2.**
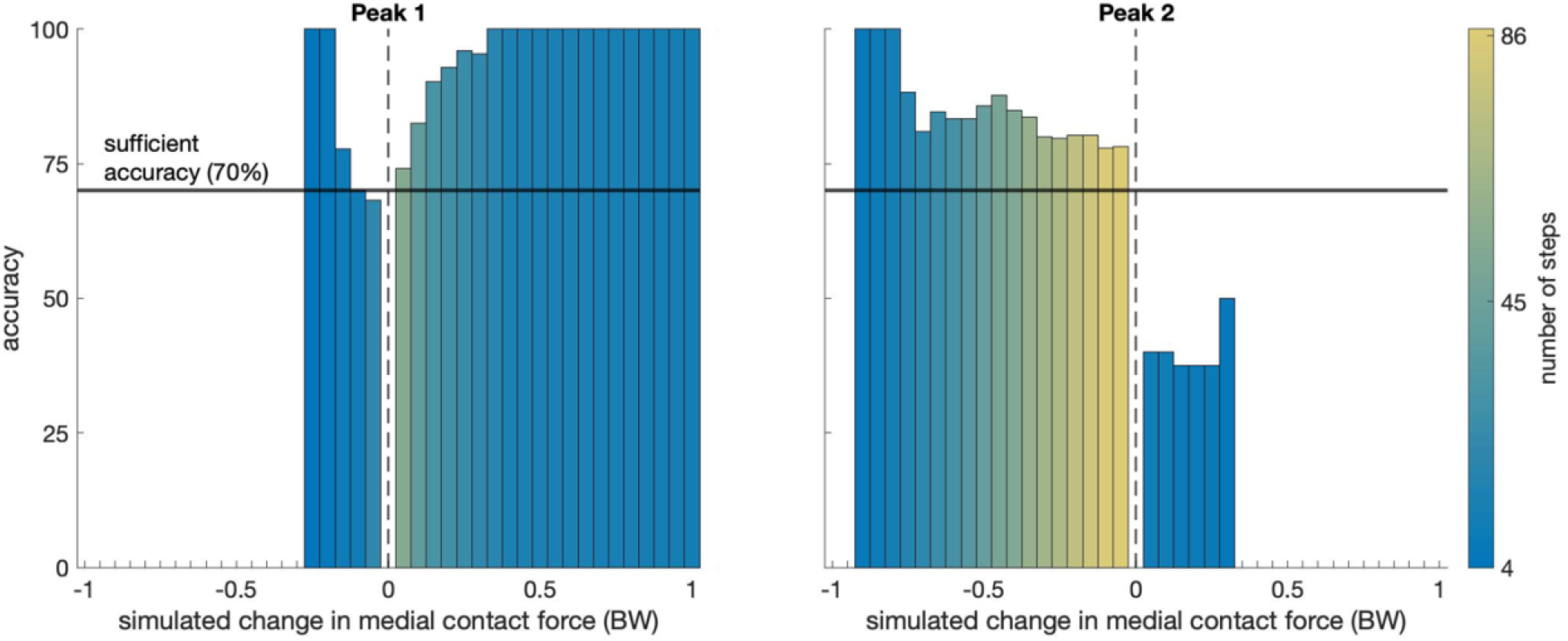
Directional accuracy (how accurately static optimization classifies an increase or decrease in medial knee contact force induced by gait modifications) versus simulated change at the first and second peaks of medial knee contact force in bodyweights (BW). Positive changes represent increases from baseline. The color of each bar denotes the number of steps at or above each simulated change threshold. For example, for all gait modification steps that static optimization predicted to be at least a 0.05 BW reduction at the first peak, about 68% of them were actually reductions, according to the instrumented knee data. Sufficient directional accuracy is 70% (horizontal black line). The smallest simulated changes in medial knee contact force that achieved sufficient directional accuracy were 0.10 BW decreases and 0.05 BW increases at the first peak, and 0.05 BW decreases at the second peak.

We evaluated directional accuracy from 0 to ±1 bodyweights (BW) in 0.05 BW increments of simulated change. If there were greater than three experimental measurements at or above the increment, we evaluated all steps for which static optimization predicted a change greater than the increment (i.e., simulated change thresholds). The directional accuracy was the proportion of these steps for which static optimization predicted the same directional change as the experimental data. We also calculated overall directional accuracies at each contact force peak for all steps (Supplementary Tables 4 & 5).

To establish a target directional accuracy for the simulations, we defined the directional accuracy of the standard of practice, which is to prescribe the same load-reducing modification to all participants (Simic et al., 2011; Hunt et al., 2018). Based on the instrumented knee data, a naïve classifier using this assumption would be at most 68% accurate (Supplementary Table 6, Supplementary Note 3). Therefore, we defined *sufficient* directional accuracy as 70% and identified the minimum simulated change threshold that achieved this accuracy for increases and decreases in both peaks of contact force estimates.

## Results

Static optimization underpredicted first peak MCF with mean absolute error of 0.16 BW, and overpredicted the second peak with mean absolute error of 0.31 BW (Fig. 1, Supplementary Table 1). The mean absolute errors of changes in MCF from baseline were 0.18 BW and 0.35 BW at the first and second peaks, respectively (Supplementary Table 2). Average stance-phase root mean square error was 0.32 BW.

Static optimization detected the directional change with sufficient accuracy (≥70%) for 0.05 BW increases at the first peak and decreases at the second peak of MCF (Fig. 2). Sufficient accuracy was also achieved for 0.10 BW decreases at the first peak. However, sufficient accuracy was not achieved for second peak increases. Overall, static optimization predicted MCF with higher directional accuracy for increases at the first peak and reductions at the second peak (Supplementary Tables 4 & 5). See Supplementary Note 2 for lateral and total knee contact force results.

## Discussion

Our objective was to determine how accurately static optimization estimates changes in peak knee MCF induced by gait modifications. We found that if static optimization detected early-stance reductions, late-stance reductions, and early-stance increases in MCF of at least 0.10 BW, it was more accurate than the naïve assumption that gait modifications affect all individuals the same way. If static optimization detects changes greater than this threshold, we can be reasonably confident that the direction of the change is correct. This finding suggests that static optimization may be useful for evaluating how gait modifications affect early-stance MCF in future studies of individuals with osteoarthritis.

Our average root mean square errors are within the ranges of past Grand Challenge winners for best predictions (MCF: 0.21-0.47 BW; lateral contact force: 0.26-0.51 BW; total contact force: 0.48-0.69 BW), who used a variety of modeling and simulation techniques, including those that incorporated deformable contact models or electromyography signals (Hast and Piazza, 2013; Jung et al., 2016; Kim et al., 2013; Kinney et al., 2013a; Knowlton et al., 2013; Manal and Buchanan, 2013; Supplementary Note 2). This suggests that a simple muscle redundancy solver has comparable error to more complex approaches for estimating knee contact force. Additionally, we confirmed that our peak total contact force errors agree with those from a similar static optimization method (Knarr and Higginson, 2015; Supplementary Note 2).

Our static optimization implementation has sufficient sensitivity for identifying clinically relevant reductions in peak total contact force and MCF derived from weight loss studies. A 7.7-10.2% BW loss has been shown to improve function and reduce knee joint loading (Atukorala et al., 2016; Messier et al., 2011). Additionally, peak total contact force reduces 1-4-fold with every unit of weight loss (Aaboe et al., 2011; DeVita et al., 2016; Messier et al., 2005). Taken together, we expect peak total contact force reductions in the range of 0.08-0.41 BW to be clinically meaningful. Our simulations detected reductions in peak total contact force at the lower end of this range (0.05-0.15 BW) with at least 70% accuracy (Supplementary Table 3). Similarly, since peak MCF comprised about 60-70% of peak total contact force in the Grand Challenge instrumented knee data, we expect that 0.05-0.29 BW peak MCF reductions are clinically meaningful. Our simulations detected peak MCF reductions at the lower end of this range (0.05-0.10 BW) with at least 70% accuracy.

It is important to identify the limitations of our study. Although a static optimization cost function that minimized sum-squared muscle activation resulted in sufficient directional accuracy for most cases (Fig. 2, Supplementary Fig. 2), this cost function may not optimally represent muscle coordination for individuals with osteoarthritis learning a new gait pattern (Shull et al., 2015). Incorporating additional cost function terms that better capture elevated cocontraction (Booij et al., 2020) or pain minimization, or using electromyography data to inform simulations (Lloyd and Besier, 2003; Pizzolato et al., 2015), might increase static optimization accuracy. Additionally, static optimization was not able to detect increases in second peak MCF with greater accuracy than a naïve approach. This may be due to the small sample size (25 steps) for second peak MCF increases (Supplementary Table 4), and the greater number of possible muscle coordination patterns in late stance (DeMers et al., 2014). Finally, since we calculated experimental MCF and lateral contact force from regression equations, they are not direct measurements. Our simulation errors may be due, in part, to this indirect comparison.

This study demonstrates that a simple modeling and simulation pipeline can detect reductions and early-stance increases in peak knee MCF from gait modifications with at least 70% accuracy. When coupled with sensing techniques for estimating kinematics and kinetics (Karatsidis et al., 2016), static optimization might be used to evaluate joint loading outside of the lab. The accessibility of these tools and our open-source simulation software (Delp et al., 2007) can enable estimates of joint contact force to be used for evaluating gait modifications in future studies and in clinical practice.

## Data availability statement

Simulation results and input data needed to replicate the simulation results in this study are available at https://simtk.org/projects/statop_val.

## CRediT authorship contribution statement

**Janelle M. Kaneda:** Conceptualization, Methodology, Software, Formal analysis, Writing – original draft, Writing – review & editing. **Kirsten A. Seagers:** Conceptualization, Methodology, Software, Writing – review & editing. **Scott D. Uhlrich:** Conceptualization, Methodology, Software, Writing – review & editing. **Julie A. Kolesar:** Software, Formal analysis, Writing – review & editing. **Kevin A. Thomas:** Methodology, Writing – review & editing. **Scott L. Delp:** Conceptualization, Resources, Supervision, Writing – review & editing.

## Letter of transmittal

All authors have made substantial contributions to all of the following: (1) the conception and design of the study or acquisition of data, or analysis and interpretation of data, (2) drafting the article or revising it critically for important intellectual content, (3) final approval of the version to be submitted. Each author has read and concurs with the content in the manuscript.

## References

Aaboe, J., Bliddal, H., Messier, S.P., Alkjær, T., Henriksen, M., 2011. Effects of an intensive weight loss program on knee joint loading in obese adults with knee osteoarthritis. Osteoarthritis and Cartilage 19, 822–828.

Ahlbäck, S., 1968. Osteoarthrosis of the knee. A radiographic investigation. Acta Radiologica: Diagnosis Suppl 277, 7–72.

Anderson, F.C., Pandy, M.G., 2001. Static and dynamic optimization solutions for gait are practically equivalent. Journal of Biomechanics 34, 153–161.

Andriacchi, T.P., Mündermann, A., Smith, R.L., Alexander, E.J., Dyrby, C.O., Koo, S., 2004. A Framework for the in Vivo Pathomechanics of Osteoarthritis at the Knee. Annals of Biomedical Engineering 32, 447–457.

Atukorala, I., Makovey, J., Lawler, L., Messier, S.P., Bennell, K., Hunter, D.J., 2016. Is There a Dose-Response Relationship Between Weight Loss and Symptom Improvement in Persons With Knee Osteoarthritis? Arthritis Care & Research 68, 1106–1114.

Booij, M.J., Richards, R., Harlaar, J., Noort, J.C. van den, 2020. Effect of walking with a modified gait on activation patterns of the knee spanning muscles in people with medial knee osteoarthritis. The Knee 27, 198–206.

Brandon, S.C.E., Miller, R.H., Thelen, D.G., Deluzio, K.J., 2014. Selective lateral muscle activation in moderate medial knee osteoarthritis subjects does not unload medial knee condyle. Journal of Biomechanics 47, 1409–1415.

Brisson, N.M., Gatti, A.A., Damm, P., Duda, G.N., Maly, M.R., 2021. Association of Machine Learning–Based Predictions of Medial Knee Contact Force With Cartilage Loss Over 2.5 Years in Knee Osteoarthritis. Arthritis & Rheumatology 73, 1638–1645.

Cui, A., Li, H., Wang, D., Zhong, J., Chen, Y., Lu, H., 2020. Global, regional prevalence, incidence and risk factors of knee osteoarthritis in population-based studies. eClinicalMedicine 29.

Delp, S.L., Anderson, F.C., Arnold, A.S., Loan, P., Habib, A., John, C.T., Guendelman, E., Thelen, D.G., 2007. OpenSim: Open-Source Software to Create and Analyze Dynamic Simulations of Movement. IEEE Transactions on Biomedical Engineering 54, 1940–1950.

DeMers, M.S., Pal, S., Delp, S.L., 2014. Changes in Tibiofemoral Forces due to Variations in Muscle Activity during Walking. Journal of Orthopaedic Research 32, 769–776.

DeVita, P., Rider, P., Hortobágyi, T., 2016. Reductions in knee joint forces with weight loss are attenuated by gait adaptations in class III obesity. Gait & Posture 45, 25–30.

D’Lima, D.D., Fregly, B.J., Patil, S., Steklov, N., Colwell, C.W., 2012. Knee joint forces: prediction, measurement, and significance. Proceedings of the Institution of Mechanical Engineers, Part H 226, 95–102.

Felson, D.T., Nevitt, M.C., Zhang, Y., Aliabadi, P., Baumer, B., Gale, D., Li, W., Yu, W., Xu, L., 2002. High prevalence of lateral knee osteoarthritis in Beijing Chinese compared with Framingham Caucasian subjects. Arthritis & Rheumatology 46, 1217–1222.

Fregly, B.J., Reinbolt, J.A., Rooney, K.L., Mitchell, K.H., Chmielewski, T.L., 2007. Design of Patient-Specific Gait Modifications for Knee Osteoarthritis Rehabilitation. IEEE Transactions on Biomedical Engineering 54, 1687–1695.

Fregly, B.J., D’Lima, D.D., Colwell, C.W., 2009. Effective Gait Patterns for Offloading the Medial Compartment of the Knee. Journal of Orthopaedic Research 27, 1016–1021.

Fregly, B.J., Besier, T.F., Lloyd, D.G., Delp, S.L., Banks, S.A., Pandy, M.G., D’Lima, D.D., 2012. Grand Challenge Competition to Predict In Vivo Knee Loads. Journal of Orthopaedic Research 30, 503–513.

Hast, M.W., Piazza, S.J., 2013. Dual-Joint Modeling for Estimation of Total Knee Replacement Contact Forces During Locomotion. Journal of Biomechanical Engineering 135.

Hicks, J.L., Uchida, T.K., Seth, A., Rajagopal, A., Delp, S.L., 2015. Is My Model Good Enough? Best Practices for Verification and Validation of Musculoskeletal Models and Simulations of Movement. Journal of Biomechanical Engineering 137, 0209051–02090524.

Hunt, M.A., Charlton, J.M., Krowchuk, N.M., Tse, C.T.F., Hatfield, G.L., 2018. Clinical and biomechanical changes following a 4-month toe-out gait modification program for people with medial knee osteoarthritis: a randomized controlled trial. Osteoarthritis and Cartilage 26, 903–911.

Jones, R.K., Chapman, G.J., Findlow, A.H., Forsythe, L., Parkes, M.J., Sultan, J., Felson, D.T., 2013. A New Approach to Prevention of Knee Osteoarthritis: Reducing Medial Load in the Contralateral Knee. The Journal of Rheumatology 40, 309–315.

Jung, Y., Phan, C.-B., Koo, S., 2016. Intra-Articular Knee Contact Force Estimation During Walking Using Force-Reaction Elements and Subject-Specific Joint Model. Journal of Biomechanical Engineering 138.

Karatsidis, A., Bellusci, G., Schepers, H.M., de Zee, M., Andersen, M.S., Veltink, P.H., 2016. Estimation of Ground Reaction Forces and Moments During Gait Using Only Inertial Motion Capture. Sensors (Basel, Switzerland) 17, 75.

Kim, Y.-H., Park, W.-M., Phuong, B.T.T., 2013. Effect of Joint Center Location on In-Vivo Joint Contact Forces During Walking. In Proceedings of the American Society of Mechanical Engineers 2010 Summer Bioengineering Conference. Naples, Florida, USA. pp. 267–268.

Kinney, A.L., Besier, T.F., D’Lima, D.D., Fregly, B.J., 2013a. Update on Grand Challenge Competition to Predict in Vivo Knee Loads. Journal of Biomechanical Engineering 135, 0210121–0210124.

Kinney, A.L., Besier, T.F., Silder, A., Delp, S.L., D’Lima, D.D., Fregly, B.J., 2013b. Changes in In Vivo Knee Contact Forces through Gait Modification. Journal of Orthopaedic Research 31, 434–440.

Knarr, B.A., Higginson, J.S., 2015. Practical approach to subject-specific estimation of knee joint contact force. Journal of Biomechanics 48, 2897–2902.

Knowlton, C.B., Wimmer, M.A., Lundberg, H.J., 2013. Grand Challenge Competition: A Parametric Numerical Model to Predict In Vivo Medial and Lateral Knee Forces in Walking Gaits. In Proceedings of the American Society of Mechanical Engineers 2012 Summer Bioengineering Conference. Fajardo, Puerto Rico, USA. pp. 199–200.

Lerner, Z.F., DeMers, M.S., Delp, S.L., Browning, R.C., 2015. How Tibiofemoral Alignment and Contact Locations Affect Predictions of Medial and Lateral Tibiofemoral Contact Forces. Journal of Biomechanics 48, 644–650.

Lloyd, D.G., Besier, T.F., 2003. An EMG-driven musculoskeletal model to estimate muscle forces and knee joint moments in vivo. Journal of Biomechanics 36, 765–776.

Lundberg, H.J., Knowlton, C., Wimmer, M.A., 2013. Fine Tuning Total Knee Replacement Contact Force Prediction Algorithms Using Blinded Model Validation. Journal of Biomechanical Engineering 135, 0210151–0210159.

Manal, K., Buchanan, T.S., 2013. Predictions of Condylar Contact During Normal and Medial Thrust Gait. In Proceedings of the American Society of Mechanical Engineers 2012 Summer Bioengineering Conference. Fajardo, Puerto Rico, USA. pp. 197–198.

Messier, S.P., Gutekunst, D.J., Davis, C., DeVita, P., 2005. Weight loss reduces knee-joint loads in overweight and obese older adults with knee osteoarthritis. Arthritis & Rheumatology 52, 2026–2032.

Messier, S.P., Legault, C., Loeser, R.F., Van Arsdale, S.J., Davis, C., Ettinger, W.H., DeVita, P., 2011. Does high weight loss in older adults with knee osteoarthritis affect bone-on-bone joint loads and muscle forces during walking? Osteoarthritis and Cartilage 19, 272–280.

Meyer, A.J., D’Lima, D.D., Banks, S.A., Coburn, J., Harman, M., Mikashima, Y., Fregly, B.J., 2013a. Evaluation of Regression Equations for Medial and Lateral Contact Force From Instrumented Knee Implant Data. In Proceedings of the American Society of Mechanical Engineers 2011 Summer Bioengineering Conference. Farmington, Pennsylvania, USA. pp. 389–390.

Meyer, A.J., D’Lima, D.D., Besier, T.F., Lloyd, D.G., Colwell, C.W., Fregly, B.J., 2013b. Are External Knee Load and EMG Measures Accurate Indicators of Internal Knee Contact Forces during Gait? Journal of Orthopaedic Research 31, 921–929.

Miyazaki, T., Wada, M., Kawahara, H., Sato, M., Baba, H., Shimada, S., 2002. Dynamic load at baseline can predict radiographic disease progression in medial compartment knee osteoarthritis. Annals of Rheumatic Diseases 61, 617–622.

Piazza, S.J., Erdemir, A., Okita, N., Cavanagh, P.R., 2004. Assessment of the functional method of hip joint center location subject to reduced range of hip motion. Journal of Biomechanics 37, 349–356.

Pizzolato, C., Lloyd, D.G., Sartori, M., Ceseracciu, E., Besier, T.F., Fregly, B.J., Reggiani, M., 2015. CEINMS: a toolbox to investigate the influence of different neural control solutions on the prediction of muscle excitation and joint moments during dynamic motor tasks. Journal of Biomechanics 48, 3929–3936.

Rajagopal, A., Dembia, C.L., DeMers, M.S., Delp, D.D., Hicks, J.L., Delp, S.L., 2016. Full body musculoskeletal model for muscle-driven simulation of human gait. IEEE Transactions on Biomedical Engineering 63, 2068–2079.

Richards, R.E., Andersen, M.S., Harlaar, J., van den Noort, J.C., 2018. Relationship between knee joint contact forces and external knee joint moments in patients with medial knee osteoarthritis: effects of gait modifications. Osteoarthritis and Cartilage 26, 1203–1214.

Seth, A., et al., 2018. OpenSim: Simulating musculoskeletal dynamics and neuromuscular control to study human and animal movement. PLOS Computational Biology 14, e1006223.

Sharma, L., Hurwitz, D.E., Thonar, E.J.-M.A., Sum, J.A., Lenz, M.E., Dunlop, D.D., Schnitzer, T.J., Kirwan-Mellis, G., Andriacchi, T.P., 1998. Knee adduction moment, serum hyaluronan level, and disease severity in medial tibiofemoral osteoarthritis. Arthritis & Rheumatism 41, 1233–1240.

Shull, P.B., Silder, A., Shultz, R., Dragoo, J.L., Besier, T.F., Delp, S.L., Cutkosky, M.R., 2013. Six-week gait retraining program reduces knee adduction moment, reduces pain, and improves function for individuals with medial compartment knee osteoarthritis. Journal of Orthopaedic Research 31, 1020–1025.

Shull, P.B., Huang, Y., Schlotman, T., Reinbolt, J.A., 2015. Muscle force modification strategies are not consistent for gait retraining to reduce the knee adduction moment in individuals with knee osteoarthritis. Journal of Biomechanics 48, 3163–3169.

Simic, M., Hinman, R.S., Wrigley, T.V., Bennell, K.L., Hunt, M.A., 2011. Gait modification strategies for altering medial knee joint load: a systematic review. Arthritis Care & Research 63, 405–426.

Uhlrich, S.D., Jackson, R.W., Seth, A., Kolesar, J.A., Delp, S.L., 2022. Muscle coordination retraining inspired by musculoskeletal simulations reduces knee contact force. Scientific Reports 12, 9842.

Vos, T., et al., 2012. Years lived with disability (YLDs) for 1160 sequelae of 289 diseases and injuries 1990-2010: a systematic analysis for the Global Burden of Disease Study 2010. Lancet 380, 2163–2196.

Walter, J.P., D’Lima, D.D., Colwell, C.W., Fregly, B.J., 2010. Decreased Knee Adduction Moment Does Not Guarantee Decreased Medial Contact Force during Gait. Journal of Orthopaedic Research 28, 1348–1354.

Zhao, D., Banks, S.A., Mitchell, K.H., D’Lima, D.D., Colwell Jr., C.W., Fregly, B.J., 2007. Correlation between the knee adduction torque and medial contact force for a variety of gait patterns. Journal of Orthopaedic Research 25, 789–797.

