## Supplementary Materials for "Can static optimization detect changes in peak medial knee contact forces induced by gait modifications?"

### Supplementary Note 1. Musculoskeletal modeling and simulation details.

We evaluated frontal-plane knee alignment and tibiofemoral contact points from a standing radiograph for the 6$^{th}$ Grand Challenge subject (Lerner et al., 2015). A radiograph was not available for the other subjects, so we determined contact points using regression equations with femoral epicondyle marker positions as inputs (Winby et al., 2009).

Non-physiologically large muscle activations for hip rotators in some trials across subjects resulted from static optimization, which led to hip-muscle coactivation. Since the primary hip rotators do not also actuate the knee, we lowered the optimizer weight on the hip rotation reserve to reduce these errant activations. We applied this modification to all simulations because it did not affect trials that did not previously have hip-muscle coactivations.

**Supplementary Note 2. Results and discussion for lateral and total knee contact forces.**

Static optimization overpredicted both peaks of lateral and total contact force, with a first peak mean absolute error of 0.81 bodyweights (BW) and 0.83 BW respectively, as well as 0.39 BW and 0.65 BW at the second peak (Supplementary Fig. 1A & 1B). The large first peak errors were primarily due to large errors from the gait modification trials, rather than the baseline trials (Supplementary Table 1). In contrast to medial contact force (MCF), the mean absolute errors for changes in lateral and total contact force from baseline were greater at the first peak than the second peak, with errors of 0.45 BW and 0.21 BW for lateral contact force, and 0.60 BW and 0.37 BW for total contact force (Supplementary Table 2). Average stance-phase root mean square error was 0.47 BW for lateral contact force and 0.66 BW for total contact force.

The minimum simulated change thresholds that achieved sufficient directional accuracy (≥70%) for lateral and total contact force were larger than those for MCF, up to 0.45 BW for total contact force first peak increases (Supplementary Fig. 2, Supplementary Table 3). The trend that static optimization predicted MCF with higher directional accuracy for increases at the first peak and reductions at the second peak was also consistent for lateral and total contact force (Supplementary Tables 4 & 5).

Although peak total contact force error was greater than peak MCF error, it is similar to those reported previously using comparable modeling and simulation methods. Our total contact force peak errors (first peak: 0.83 BW, second peak: 0.65 BW) and average stance-phase root mean square error (0.66 BW) are within ranges reported by Knarr and Higginson (first peak: 0.14-0.95 BW; second peak: 0.40-1.37 BW; root mean square error: 0.37-0.67 BW), who used a default static optimization implementation in OpenSim and simulated for the first three Grand Challenge subjects (2015).

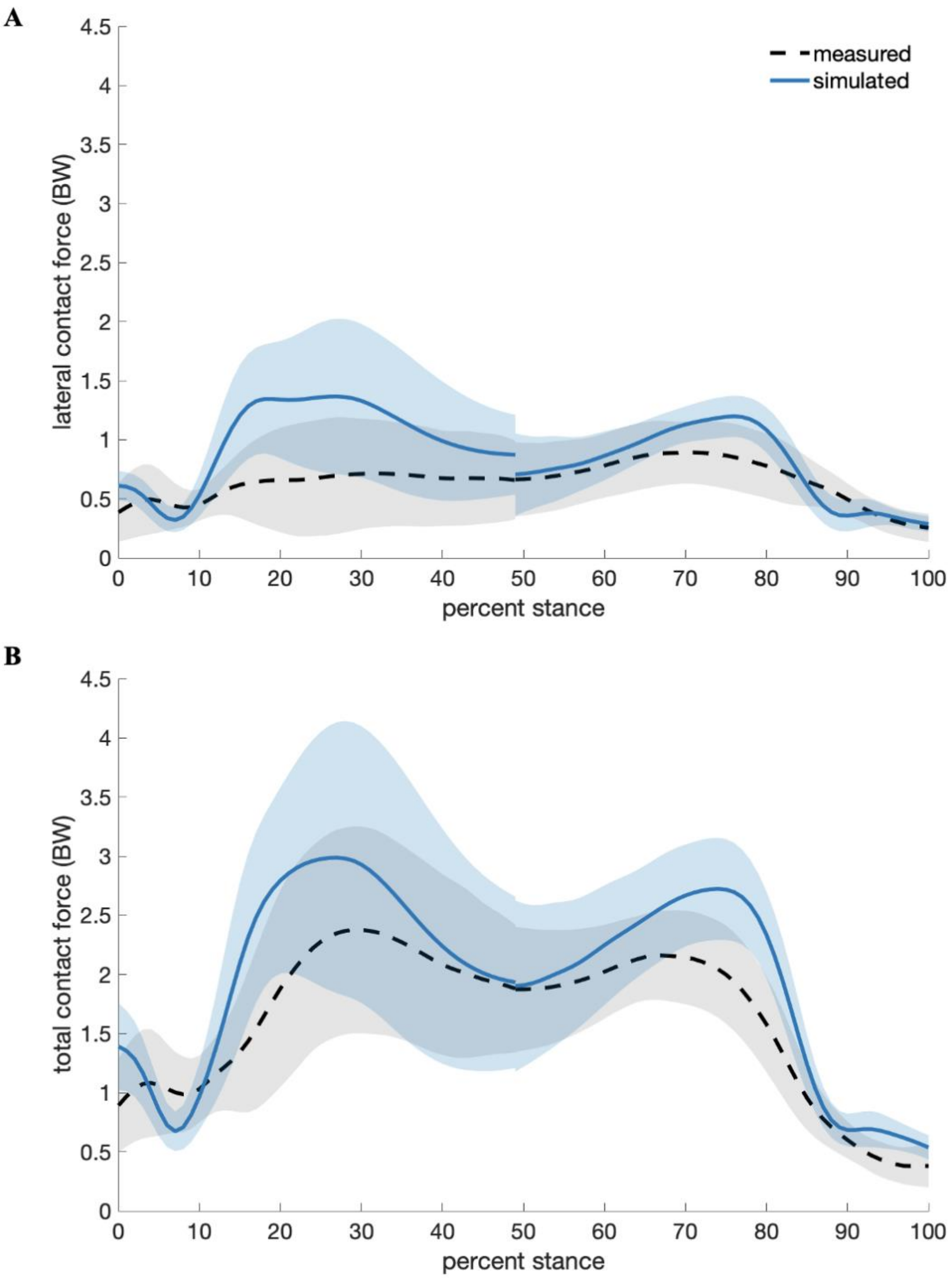


**Supplementary Figure 1.** Mean ± standard deviation (shaded) of measured and simulated lateral (**A**) and total (**B**) knee contact force in bodyweights (BW) for all evaluated trials, including natural walking (*N*=115). The discontinuity at 50% of stance results from computing early-stance and late-stance contact force from different gait cycles for some steps.

**Supplementary Table 1.** Mean absolute errors (in bodyweights, BW) for both peaks of medial, lateral, and total knee contact force (MCF, LCF, TCF) per walking pattern, listed in order of decreasing sample size (*n*).

| Walking pattern (*n*) | MCF Peak 1 (BW) | MCF Peak 2 (BW) | LCF Peak 1 (BW) | LCF Peak 2 (BW) | TCF Peak 1 (BW) | TCF Peak 2 (BW) |
|---|---|---|---|---|---|---|
| **Walking poles (31)** | 0.09 | 0.19 | 0.92 | 0.41 | 0.88 | 0.56 |
| **Bouncy (16)** | 0.15 | 0.39 | 1.10 | 0.42 | 1.22 | 0.78 |
| **Crouch (15)** | 0.21 | 0.28 | 1.09 | 0.34 | 1.24 | 0.67 |
| **Forefoot strike (15)** | 0.24 | 0.38 | 0.75 | 0.41 | 0.84 | 0.79 |
| **Medial thrust (13)** | 0.13 | 0.26 | 0.88 | 0.32 | 0.77 | 0.51 |
| **Smooth (7)** | 0.29 | 0.29 | 0.18 | 0.37 | 0.42 | 0.60 |
| **Trunk sway (4)** | 0.20 | 0.42 | 0.37 | 0.28 | 0.16 | 0.73 |
| **Baseline (14)** | 0.13 | 0.47 | 0.33 | 0.44 | 0.31 | 0.71 |
| **All patterns (115)** | 0.16 | 0.31 | 0.81 | 0.39 | 0.83 | 0.65 |

**Supplementary Table 2.** Mean absolute errors (in bodyweights, BW) of changes from baseline, for both peaks of medial, lateral, and total knee contact force (MCF, LCF, TCF). Changes were calculated by subtracting the average baseline peak from each gait modification trial peak for each subject. *N*=101 gait modification trials.

| | **Peak 1 (BW)** | **Peak 2 (BW)** |
|---|---|---|
| **Medial contact force** | 0.18 | 0.35 |
| **Lateral contact force** | 0.45 | 0.21 |
| **Total contact force** | 0.60 | 0.37 |

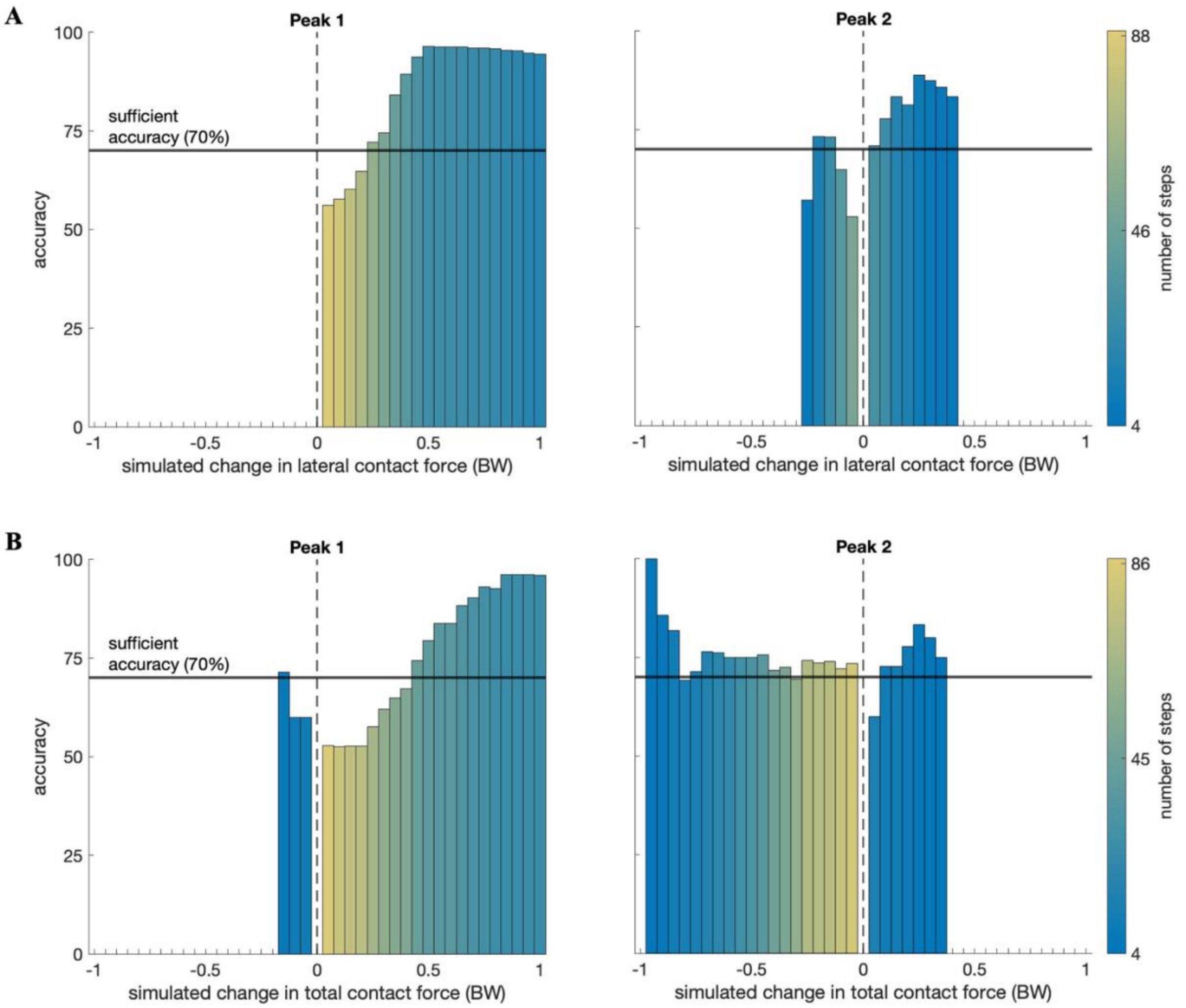


**Supplementary Figure 2.** Directional accuracy (how accurately static optimization classifies an increase or decrease in knee contact force induced by gait modifications) versus simulated change at the first and second peaks of lateral (**A**) and total (**B**) knee contact force in bodyweights (BW). Positive changes represent increases from baseline. The color of each bar denotes the number of steps at or above each simulated change threshold. For example, for all gait modification steps that static optimization predicted to be at least a 0.05 BW increase at Peak 1 of lateral knee contact force (**A**, left), about 56% of them were actually increases, according to the instrumented knee data. Sufficient directional accuracy is 70% (horizontal black line). The smallest simulated changes in lateral and total knee contact force that achieved sufficient directional accuracy are reported in Supplementary Table 3.

**Supplementary Table 3.** The smallest simulated changes in lateral and total knee contact force that achieved sufficient directional accuracy (≥70%), reported in bodyweights (BW).

| | | **Lateral contact force (BW)** | **Total contact force (BW)** |
|---|---|---|---|
| Peak 1 | Increase | 0.25 | 0.45 |
| | Decrease | N.A.* | 0.15 |
| Peak 2 | Increase | 0.05 | 0.10 |
| | Decrease | 0.15 | 0.05 |

* Not Applicable. No viable data points.

**Supplementary Table 4.** Proportions of experimental peak increases that were correctly predicted for medial, lateral, and total knee contact force (MCF, LCF, TCF) per gait modification, listed in order of decreasing total sample size (*n*).

| Gait modification (*n*) | MCF Peak 1 (%) | MCF Peak 2 (%) | LCF Peak 1 (%) | LCF Peak 2 (%) | TCF Peak 1 (%) | TCF Peak 2 (%) |
|---|---|---|---|---|---|---|
| **Walking poles (31)** | 8 / 10 (80%) | 0 / 1 (0%) | 12 /12 (100%) | 0 / 7 (0%) | 6 / 6 (100%) | 0 / 3 (0%) |
| **Bouncy (16)** | 14 / 14 (100%) | 1 / 4 (25%) | 16 / 16 (100%) | 6 / 7 (85.7%) | 14 / 14 (100%) | 1 / 3 (33.3%) |
| **Crouch (15)** | 10 / 12 (83.3%) | 2 / 4 (50%) | 8 / 8 (100%) | 7 / 9 (77.8%) | 11 / 11 (100%) | 3 / 5 (60%) |
| **Forefoot strike (15)** | 3 / 8 (37.5%) | 1 / 6 (16.7%) | 9 / 13 (69.2%) | 2 / 9 (22.2%) | 6 / 9 (66.7%) | 3 / 7 (42.9%) |
| **Medial thrust (13)** | 6 / 7 (85.7%) | 0 / 7 (0%) | 5 / 5 (100%) | 4 / 7 (57.1%) | 6 / 7 (85.7%) | 1 / 7 (14.3%) |
| **Smooth (7)** | 2 / 3 (66.7%) | 1 / 3 (33.3%) | 2 / 2 (100%) | 2 / 7 (28.6%) | 3 / 3 (100%) | 0 / 4 (0%) |
| **Trunk sway (4)** | 1 / 4 (25%) | N.A.* | 1 / 1 (100%) | 3 / 4 (75%) | 2 / 3 (66.7%) | 1 / 2 (50%) |
| **All modifications (101)** | 44 / 58 (75.9%) | 5 / 25 (20%) | 53 / 57 (93%) | 24 / 50 (48%) | 48 / 53 (90.6%) | 9 / 31 (29%) |

* Not Applicable. Zero experimental changes at the given knee contact force and peak type.

**Supplementary Table 5.** Proportions of experimental peak reductions that were correctly predicted for medial, lateral, and total knee contact force (MCF, LCF, TCF) per gait modification, listed in order of decreasing total sample size (*n*).

| Gait modification (*n*) | MCF Peak 1 (%) | MCF Peak 2 (%) | LCF Peak 1 (%) | LCF Peak 2 (%) | TCF Peak 1 (%) | TCF Peak 2 (%) |
|---|---|---|---|---|---|---|
| **Walking poles (31)** | 10 / 21 (47.6%) | 30 / 30 (100%) | 0 / 19 (0%) | 15 / 24 (62.5%) | 2 / 25 (8%) | 28 / 28 (100%) |
| **Bouncy (16)** | 0 / 2 (0%) | 12 / 12 (100%) | N.A.* | 6 / 9 (66.7%) | 0 / 2 (0%) | 11 / 13 (84.6%) |
| **Crouch (15)** | 2 / 3 (66.7%) | 11 / 11 (100%) | 0 / 7 (0%) | 2 / 6 (33.3%) | 0 / 4 (0%) | 10 / 10 (100%) |
| **Forefoot strike (15)** | 5 / 7 (71.4%) | 5 / 9 (55.6%) | 0 / 2 (0%) | 6 / 6 (100%) | 3 / 6 (50%) | 6 / 8 (75%) |
| **Medial thrust (13)** | 3 / 6 (50%) | 5 / 6 (83.3%) | 4 / 8 (50%) | 4 / 6 (66.7%) | 1 / 6 (16.7%) | 6 / 6 (100%) |
| **Smooth (7)** | 0 / 4 (0%) | 3 / 4 (75%) | 1 / 5 (20%) | N.A.* | 1 / 4 (25%) | 1 / 3 (33.3%) |
| **Trunk sway (4)** | N.A.* | 4 / 4 (100%) | 0 / 3 (0%) | N.A.* | 0 / 1 (0%) | 2 / 2 (100%) |
| **All modifications (101)** | 20 / 43 (46.5%) | 70 / 76 (92.1%) | 5 / 44 (11.4%) | 33 / 51 (64.7%) | 7 / 48 (14.6%) | 64 / 70 (91.4%) |

* Not Applicable. Zero experimental changes at the given knee contact force and peak type.

**Supplementary Note 3. Determining sufficient directional accuracy based on the standard of practice.**

To determine the directional accuracy of the standard of practice, we used the changes at stance-phase peaks of the instrumented knee replacement (IKR) force data. The current standard of practice for assigning gait modifications is a non-personalized approach that assumes that a given modification has the same effect on all individuals (Simic et al., 2011; Hunt et al., 2018). Based on previous literature (Fregly et al., 2009; Kinney et al., 2013; Razu and Guess, 2018; Simic et al., 2012; Steele et al., 2012), we assumed that all gait modifications used in our study decreased both peaks of medial, lateral, and total knee contact force, except for the first peak of bouncy gait and both peaks of crouch gait, which were assumed to increase.

For each IKR gait modification trial, we calculated the peak directional change the same way as the present study, that is, subtracting the average IKR baseline peak from each IKR gait modification trial peak. We calculated directional accuracy as the proportion of peak directional changes that matched the assumed direction from literature (Supplementary Table 6). To identify the minimum simulated change thresholds that surpass the maximum directional accuracy determined from experimental data alone, we defined sufficient directional accuracy as 70%, which is 2% greater than the maximum IKR-based directional accuracy (medial contact force, Peak 2).

**Supplementary Table 6.** Directional accuracies at the first and second peaks of knee contact forces calculated solely from the instrumented knee data, assuming the same target directional changes across all subjects from each gait modification.

| | **Peak 1** | **Peak 2** |
|---|---|---|
| **Medial contact force** | 0.63 | 0.68 |
| **Lateral contact force** | 0.60 | 0.53 |
| **Total contact force** | 0.66 | 0.64 |

### Supplementary References


Fregly, B.J., D'Lima, D.D., Colwell, C.W., 2009. Effective Gait Patterns for Offloading the Medial Compartment of the Knee. Journal of Orthopedic Research 27, 1016–1021.

Hunt, M.A., Charlton, J.M., Krowchuk, N.M., Tse, C.T.F., Hatfield, G.L., 2018. Clinical and biomechanical changes following a 4-month toe-out gait modification program for people with medial knee osteoarthritis: a randomized controlled trial. Osteoarthritis and Cartilage 26, 903–911.

Kinney, A.L., Besier, T.F., Silder, A., Delp, S.L., D'Lima, D.D., Fregly, B.J., 2013. Changes in In Vivo Knee Contact Forces through Gait Modification. Journal of Orthopaedic Research 31, 434–440.

Knarr, B.A., Higginson, J.S., 2015. Practical approach to subject-specific estimation of knee joint contact force. Journal of Biomechanics 48, 2897–2902.

Lerner, Z.F., DeMers, M.S., Delp, S.L., Browning, R.C., 2015. How Tibiofemoral Alignment and Contact Locations Affect Predictions of Medial and Lateral Tibiofemoral Contact Forces. Journal of Biomechanics 48, 644–650.

Razu, S.S., Guess, T.M., 2018. Electromyography-Driven Forward Dynamics Simulation to Estimate In Vivo Joint Contact Forces During Normal, Smooth, and Bouncy Gaits. Journal of Biomechanical Engineering 140, 0710121–0710128.

Simic, M., Hinman, R.S., Wrigley, T.V., Bennell, K.L., Hunt, M.A., 2011. Gait modification strategies for altering medial knee joint load: a systematic review. Arthritis Care & Research 63, 405–426.

Simic, M., Hunt, M.A., Bennell, K.L., Hinman, R.S., Wrigley, T.V., 2012. Trunk lean gait modification and knee joint load in people with medial knee osteoarthritis: the effect of varying trunk lean angles. Arthritis Care & Research 64, 1545–1553.

Steele, K.M., DeMers, M.S., Schwartz, M.S., Delp, S.L., 2012. Compressive Tibiofemoral Force during Crouch Gait. Gait & Posture 35, 556–560.

Winby, C.R., Lloyd, D.G., Besier, T.F., Kirk, T.B., 2009. Muscle and external load contribution to knee joint contact loads during normal gait. Journal of Biomechanics 42, 2294–2300.
